# FiCOPS: Hardware/Software Co-Design of FPGA Computational Framework for Mass Spectrometry-Based Peptide Database Search

**DOI:** 10.64898/2026.02.15.706012

**Authors:** Sumesh Kumar, Joseph Zambreno, Ashfaq Khokhar, Shoaib Akram, Fahad Saeed

**Author notes:** Corresponding author: Fahad Saeed.

## Abstract

Improving the speed and efficiency of database search algorithms that deduce peptides from mass spectrometry (MS) data has been an active area of research for more than three decades. The significance of the need for faster database search methods has rapidly increased due to the growing interest in studying non-model organisms, meta-proteomics, and proteogenomic data, which are notorious for their enormous search space. Poor scalability of serial algorithms with the growing size of the database and increasing parameters of post-translational modifications is a widely recognized problem. While high-performance computing techniques can be used on supercomputing machines, the need for real-time, on-the-instrument solutions necessitates the development of an efficient sytem-on-chip that optimizes design constraints such as cost, performance, and power of the system. To show case that such a system can work, we present an FPGA-based computational framework called FiCOPS to accelerate database search using a hardware/software co-design methodology. First, we theoretically analyze the database-search algorithm (closed-search) to reveal opportunities for parallelism and uncover computational bottlenecks. We then design an FPGA-based architectural template to exploit parallelism inherent in the search workload. We also formulate an analytical performance model for the architecture template to perform rapid design space exploration and find a near-optimal accelerator configuration. Finally, we implement our design on the Intel Stratix 10 FPGA platform and evaluate it using real-world datasets. Our experiments demonstrate that FiCOPS achieves 3.5 × speed-up over existing CPU solutions and 3× and 5× reduction in power consumption compared to existing CPU and GPU solutions.

## I. Introduction

**M**ASS spectrometry is the foundation of high-throughput proteomics, meta-proteomics, and proteogenomic studies [1]. For the past 30 years, researchers have tried to improve the efficiency of database search algorithms that deduce peptides from Mass Spectrometry (MS) data. The significance of the need for faster database search methods has rapidly increased due to the growing interest in studying nonmodel organisms, proteogenomics, and meta-proteomics that can result in traversing an enormous search space. Traditional serial algorithms [2] exhibit poor scalability with increasing size of theoretical search space (such as open-search) [3]. While database search pipelines are ubiquitous, these tools can result in a cascade of inaccuracies that are well known and prevalent including misidentification/no-identification of peptides, inconsistencies between search engines, and the tendency of the pipelines to identify most abundant peptides [4]. A vast diversity of post-translational modifications (PTMs) is unaccounted for in typical MS database search, and remains unidentified [3] due to various heuristics used to scale the search process. In the recent decade, advances in high performance architecture have provided a critical step, making it possible to develop more accurate and scalable pipelines for MS data analysis which could outperform traditional serial methods. Incorporating proteomics profiling in a clinical setting is an active goal of systems biology researchers, and such rapid analysis techniques may have application in personalized and precision medicine in identifying disease biomarkers, treatment and prognosis, and understanding of disease pathology. Recent advancements in mass spectrometry (MS) instrumentation have significantly enhanced the quality of spectra obtained from complex clinical samples, enabling more reliable discovery of clinically relevant biomarkers.

In a typical database-search, experimentally generated highdimensional and noisy MS data (called spectra) are compared against a database of protein sequences with the goal of assigning the correct sub-sequence (peptide sequence) to each spectrum [5]. Depending on the *a priori* PTM settings, the subsequences are converted to theoretical spectra used in the comparisons. Since *a priori* PTM parameters result in a theoretical database which may be exponentially larger than the original database, various filtering mechanisms are employed to reduce the search space at the expense of accuracy and specificity. Despite these filtering mechanisms, the theoretical spectra database can easily reach terabyte volumes [2], [6]. Previous studies indicate a decreased scalability of the algorithms [6] with increasing size of the database due to excessive paging and data-movement [7]. Given that many of the serial algorithms have resorted to using filtering mechanism [8], and/or can only work with a limited number of *a priori* modifications have resulted in studies of more abundant peptides/proteins, thus contributing to the bias problem [3]. Multiple recent investigations [9]–[11] aimed at the understudied proteins further illustrate the urgency to address this bias. Possible solutions include performing an unrestricted search by relaxing the mass filter or incorporating all possible PTMs in the search process when constructing the peptide database.

However, an unconstrained search and inclusion of PTMs results in a combinatorial increase in the database size leading to a gigantic search space that requires unjustifiable time and computational resources [6], [12]. The current state-of-the-art database search frameworks lead to impractical search times (several days to weeks) when searching for a large number of PTMs [6]. Therefore, novel and more efficient techniques that can traverse the MS data search-space efficiently are urgently needed. None of the existing methods cater to real-time on-the-instrument computations, and therefore, require substantial research and development efforts.

In this paper, we present a hardware/software co-design implemented on a CPU-FPGA architecture, called *FiCOPS*, to efficiently accelerate protein database search. Using “base” architectural components that would constitute a database-search engine, we develop an analytical performance model that can predict resource utilization and enable rapid design space exploration (DSE). The DSE problem is then formulated as an optimization problem to minimize search time, resource utilization, or communication costs - and demonstrate the feasibility of the design. The main components of FiCOPS design yield a workable parameterized architectural template which is then used for all experimentation and scalability results. The implementation was carried out on an Intel Stratix 10 FPGA and was evaluated extensively with benchmark datasets for a range of search parameters.

Our comprehensive experimentation shows that FiCOPS outperforms several existing serial and parallel database peptide search tools while producing correct and consistent peptide identifications. In closed search experiments, FiCOPS is 101× faster than X!Tandem [13], 82× faster than Crux [14], 3× faster than MSFragger [15] and 16× faster than GPU-Tide [16]. In open search experiments, FiCOPS achieves 4× speedup against MSFragger, 2× speed-up against HICOPS [6], and 7× speed-up against GPU-Tide. Note that like HiCOPS, FiCOPS does not propose a new database search algorithm and instead relies on underlying search-engine algorithmic workflow for peptide identification accuracy. Finally, we conduct extensive performance evaluation and report that FiCOPS consumes 32.768W of average power compared to 106W consumed by MSFragger and 132W consumed by GPU Tide – collectively depicting a highly scalable and resource-efficient parallel performance suitable for a system-on-chip (SoC) design.

## II. Background

### A. Mass Spectra Generation for Bottom-up Omics

A high-level overview of the MS-based peptide identification in bottom-up or shotgun proteomics is illustrated in Fig. 1. A biological sample containing a complex mixture of proteins is digested by an enzyme (usually Trypsin) which breaks down the protein in smaller pieces. The subsequent separation of the resulting (smaller) peptides is accomplished by liquid chromatography (LC) which are then fragmented when they elute from the LC column. This then results in the MS data, which is again scanned, and fragmented to get a MS/MS spectrum. The fragment ions in the MS/MS spectra contains the “signature” that can be used for peptide identification. From the data science prospective, each of the MS/MS spectra (hereafter *spectra*), contains the mass to charge (m/z) value of every fragment ion found in the peptide, and the associated intensity of that ion (which depicts the abundance of that specific ion). This cycle can be repeated till all the peptides have elutded from the chromatography column and can result in millions of spectra from a single experiment [17]. While spectra have all the information, the floating-point two-dimensional data needs to be translated into sequences of amino acids, called *peptides*. The most common computational tool used to make this peptide deduction are called ‘database-search’ algorithms.

**Fig. 1.**
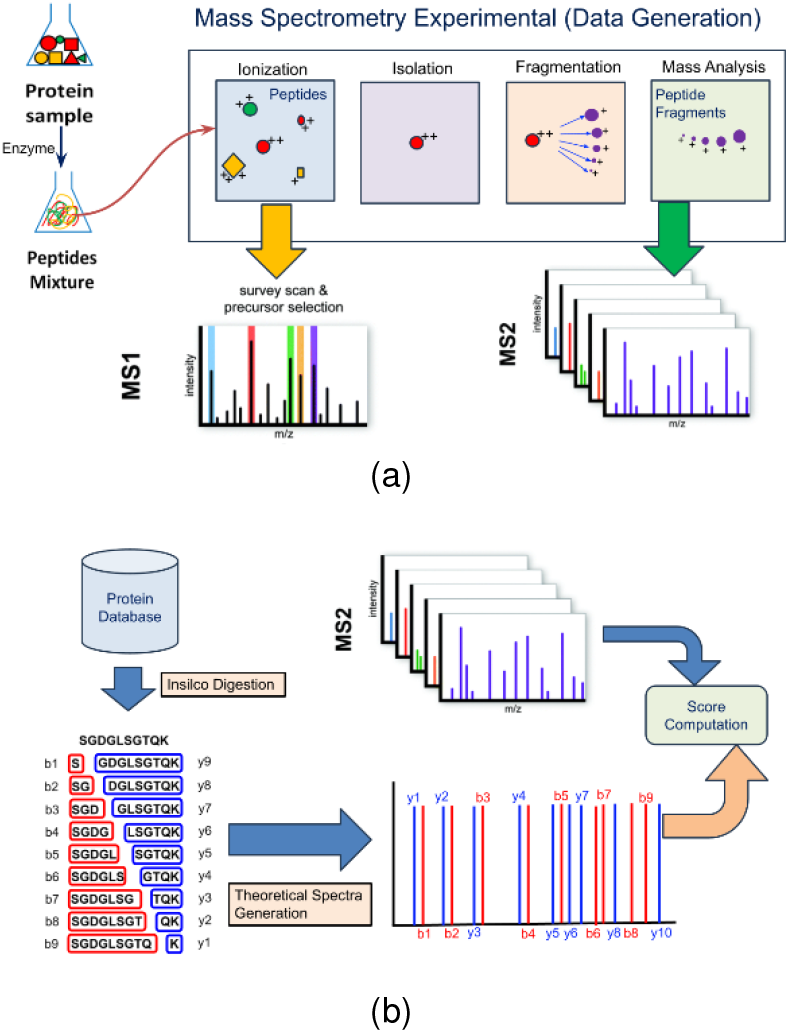
(a) Generation of MS/MS experimental spectra (b) A list of peptides is generated by an in-silico digestion of protein database. Each candidate peptide is converted into a theoretical spectra and compared with experimental spectra by computing a similarity score

### B. Database Search Algorithms

#### 1) Peptide database and theoretical spectra generation

This step begins by an *in-silico* digestion of protein sequence database by breaking down the long chains of amino acids at specific cleavage sites to generate a list of non-redundant peptides, called protein database (hereafter: *database*). This database is used to calculate all putative peptide candidates for a given set of parameters usually in the form of proteolytic enzymes, mis-cleavages, and posttranslational modifications. These subsets of peptide sequences are then used to generate a fragmentation pattern (with b- y- and other ions) such that each theoretical peptide generates a single theoretical spectrum. The addition of post-translation modification (PTM) results in addition of weights (from the modification) which results in shifting of the spectra or reduction of the peaks depending on the kind of PTM being considered. This process therefore injects newer versions of the spectra, called modified spectra, and increase the size of the database by a factor of 2^*N*^, where *N* is the number of modifications considered in the search process. The theoretical peptide generated are called *candidate peptides*, and the spectra that is theoretically generated are usually called *candidate or theoretical spectra*. The theoretical spectrum is generated using the method described in [6] by following eq. 1, where *b*[*n*] is *nth* type-b ion, *y*[*n*] is *nth* type-y ion, *A*[*n*] is the mass of *nth* amino acid in the peptide, and M is peptide mass.

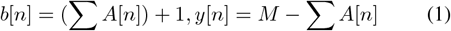

This step predicts the m/z values of all *b* and *y* ion peaks in the theoretical spectrum for a given peptide, and assigns a constant intensity value of 50 to all the peaks. The spectra for which deduction needs to be computed is called *experimental spectra*.

#### 2) Database search using peptide-indexing

To deduce a peptide for an experimental spectrum, *candidate peptides* are filtered based on their mass from the peptide database. For example, if the precursor mass of experimental spectrum is 200 Daltons (Da) and the mass filter window size is 10 Da, then all the peptides which have masses between 195 Da and 205 Da would be the *candidate peptides* for which theoretical spectra is generated, and then compared against the *experimental spectrum*.

##### Algorithm 1

Database search comparison without indexing

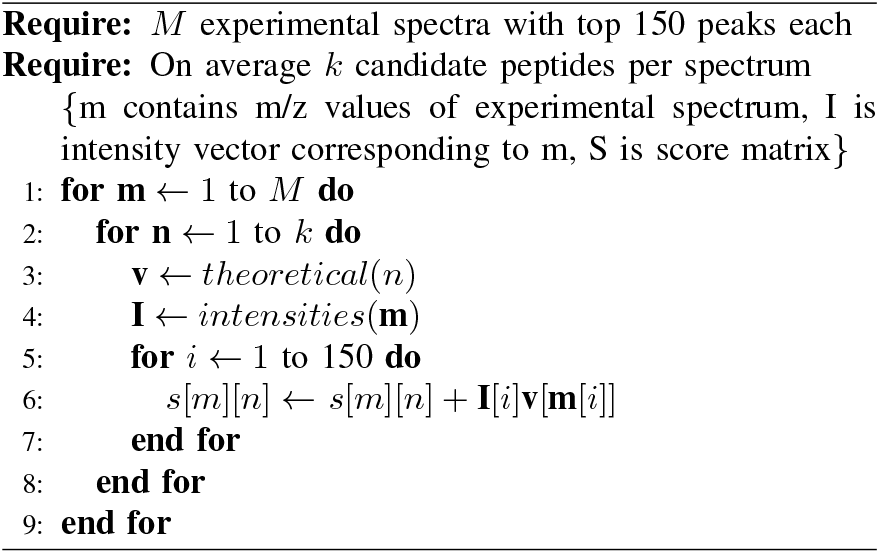

Comparison is performed using *dot-product* that is computed between the candidate spectra, and experimental spectra to quantify the similarity between them. To compute the dotproduct, the experimental and theoretical spectra are treated as arrays, where m/z is an index and intensity is the value at the corresponding index. A pseudocode illustrating the database search computation is shown in Algorithm 1. In this algorithm, for each experimental spectrum *m*, a set of *k* candidate peptides are filtered from the database based on the mass of the spectrum. For each candidate peptide *n*, a sparse *theoretical* spectrum *v* is generated, where *v* is an array whose index corresponds to the predicted m/z peak and value corresponds to the constant intensity e.g., a peptide with predicted peaks at m/z values of 147, 331, 430 will generate a sparse array containing the value 50 at only these three indicesand all the other indices will be zero. In the innermost loop, peak matching is performed with 150 peaks by searching for each m/z value of experimental spectrum in the sparse array *v*. Top 150 intensity-value peaks are selected based on empirical evidence in the field [18], [19]. After computation, candidate peptides are sorted by their score, and only top matches are reported.

#### 3) Database search using fragment-ion indexing

In a typical experiment, similarity score computation is the most time-consuming step in database-search because it requires computation of a dot product between two sparse vectors by performing the expensive inner-join step. A naive implementation of inner-join process leads to quadratic time complexity *O*(*mn*), where *m* and *n* are non-zero elements in experimental vector and theoretical vector, respectively. In general, to perform the inner-join in *O*(*n*) time, the sparse experimental vector is expanded in memory, and inner-join is performed by using array lookup for each theoretical ion, as shown in *Algorithm 1*. However, this approach still performs redundant computations because the experimental vector is highly sparse (sparsity ratio ¡ 0.05), and majority of the array lookups do not contribute much to the score.

Computationally improving this score calculation is accomplished by creating a *fragment-ion index* from the database and then searching the experimental spectra against that index [15]. In this method, the (b- and y-ion) ions are predicted to generate theoretical spectra for all the peptides. Next, the theoretical spectra are rearranged as ¡key, value¿ pairs in the form of a dictionary as shown in Fig. 2. A *key* is created corresponding to each fragment-ion m/z, and the *value* contains the list of all the peptides that had the fragment-ion (stored in the *key*). For each key, the list of peptides is stored by their molecular mass to facilitate quick retrieval of candidates. Although this process alleviates the redundant computations, it increases the memory and disk space requirements by up to 50× due to duplication of peptides references.

**Fig. 2.**
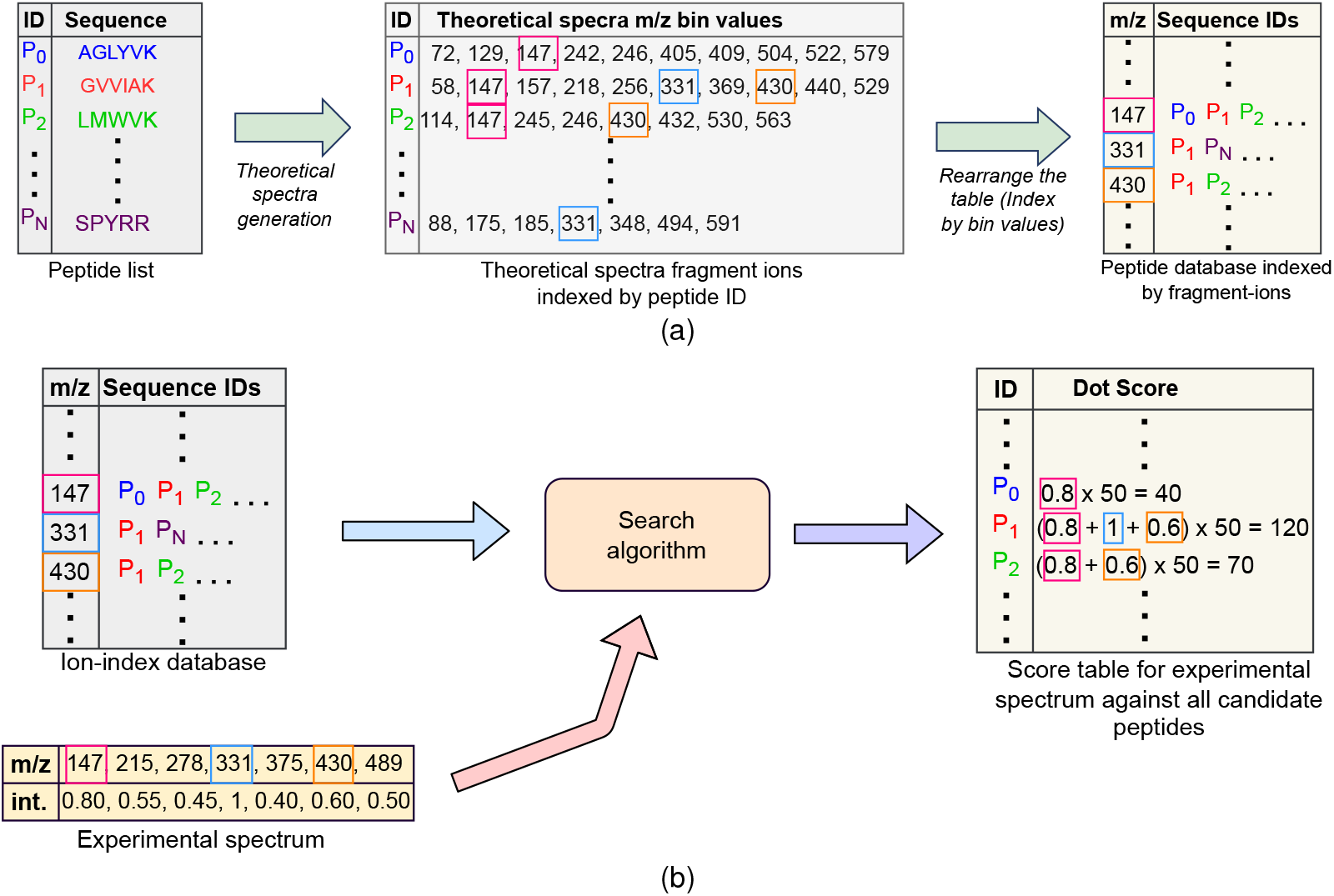
(a) Fragment-ion indexed database generation (b) Searching an experimental spectrum against ion-indexed database

##### Algorithm 2

Compute with fragment-ion index

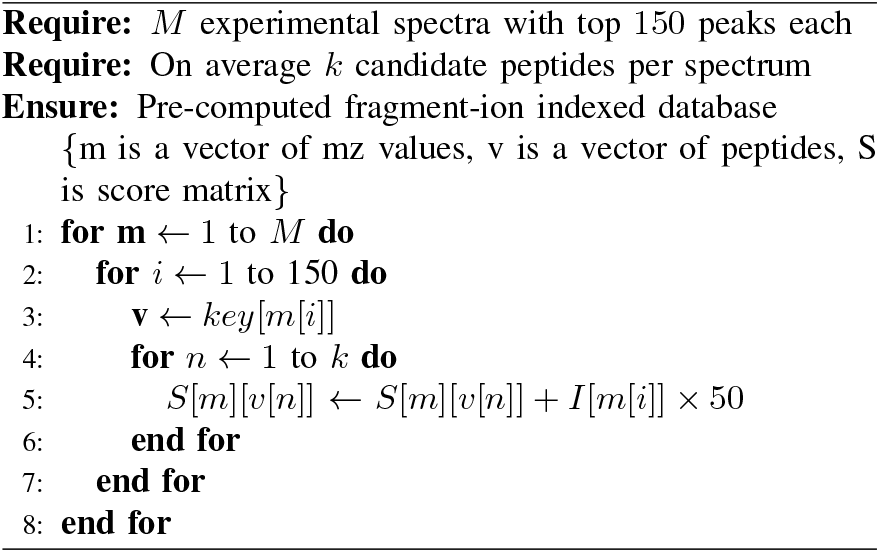

Algorithm 2 shows the pseudocode to search experimental spectra against a fragment-ion index database. In this method, for each experimental spectrum vector *m*, every ion peak is matched against all the candidate peptides. Thus, in the middle loop, for each ion peak *i*, candidate peptides *v* is retrieved from the index using the m/z value of the ion as the key. In the inner loop, the score between an experimental spectrum, and a candidate peptide is partially updated in each iteration by accumulating the product of intensity of experimental ion and a constant intensity 50 assigned to the theoretical fragment ions.

##### Remarks

Most popular database search frameworks that run on commodity computers SEQUEST [20], Crux [14], Comet [21], X!tandem [13], and MSFragger [15] have tried to solve the scalability problem by developing indexing techniques to improve the runtimes with different tradeoffs. Two indexing techniques: (1) peptide index that groups the peptides in the database by their molecular mass and use that for any further search, (2) fragment-ion index that first generates all the fragment-ions, and then groups the peptides by the shared fragment-ion mass and use the fragment-ion index as a way to accomplish any further search, especially for open-searches. Both techniques have tradeoffs with our results showing that fragment-ion indexing improves the search time by ten fold, while consuming 50× more disk space. Most of these serial algorithms do not exploit HPC capabilities of the underlying hardware. With the ubiquitous heterogenous architectures now available in the form of multicore and GPUs in large supercomputers, there is a missed opportunity that must be seized for MS analysis. Some related work on parallelization of database search methods is detailed below.

## III. SYSTEM DESIGN

FICOPS architecture design require two components that are essential for scalable processing. The first is the source of parallelism that can be exploited for Algorithm 1 and Algorithm 2. The second is the parallel architecture design template that can used to exploit the said parallelism for a given hardware. The third part of the design is the choice of parameters that are used for the designed template to run the experiments for scalability. We design all three steps that were executed here which resulted in a scalable design.

### A. Step1: Sources of Exploitable Parallelism

Carefully considering the source of parallelism inherent in these algorithms, and exploiting such source of parallelism for a given hardware could help in accelerating the search process. Algorithm 1 and Algorithm 2 shows the pseudocode for database search with peptide-index and fragment-ion index, respectively. While the overall computation can be categorized for both algorithms by three nested for-loops – not all components are parallelizable for both methods.

#### Inner loop

In Algorithm 1, lines 4-7 compute the dotproduct score between an experimental vector and a theoretical vector by iterating over 150 experimental spectrum peaks, and performing the inner-join operation. This step can be parallelized by unrolling the loop on a vector processing hardware. In Algorithm 2, however, the inner loop updates the score between experimental spectrum, and all candidate peptides. This loop execution cannot be unrolled, because each iteration performs a memory-write operation at random locations in the score matrix.

#### Middle loop

For both Algorithm 1, and Algorithm 2, the middle loop reads the candidate peptides from the memory. There is no data-dependency between the iterations of the loop, and iterations can be executed in parallel. Since each iteration performs memory read operation, the amount of exploitable parallelism is limited by memory bandwidth.

#### Outer loop

In both Algorithm 1 and Algorithm 2, the outer loop processes a single spectrum in each iteration. While there is no data-dependency for processing each experimental spectra, the peptide or fragment-ion database is stored in the shared memory which can be a potential bottleneck. Coarsegrained parallelism is still possible by utilizing multicore processors where each core operates on a separate spectrum. Multiprocessing may also be exploited when each processor has a dedicated main memory to alleviate memory bandwidth bottleneck.

### B. Step 2: Proposed FICOPS System Design: Parallel Architectural Template

Based on the parallelism analysis in step 1, we designed an architecture template for implementation of database search. Although fragment-ion indexes are popular in serial programs, our analysis reveals that their memory-intensive nature is not suitable for memory-constrained FPGA and SoC devices. Therefore, we made the design decision to develop the database search with peptide-indexing algorithm on the FPGA. While peptide-indexing loses some of the parallelism granularity, our novel design exploits *loop-unrolling* of dotproduct computation. Our designed and developed system is a configurable, parameterized, multi-core processing architecture, where the parameters control the course, and fine-grained parallelism exploited in the architecture.

### 1) High-level system architecture

The overall high-level systems architecture is shown in Fig. 3. To achieve significant speedups, the system instantiates multiple processing units (PU) that operate on distinct groups of experimental spectra. Each PU in the system contains control registers which are exposed to the CPU host for communication via the PCIe bus. This enables the CPU host to copy batches of experimental spectra and candidate peptides from main memory to each PU’s on-chip RAM, directly, via the PCIe link in a round-robin fashion. The host program polls to check if there is an update to the results, or if new data is required for each pass. Below we will discuss the design of each of the components in FICOPS architecture.

**Fig. 3.**
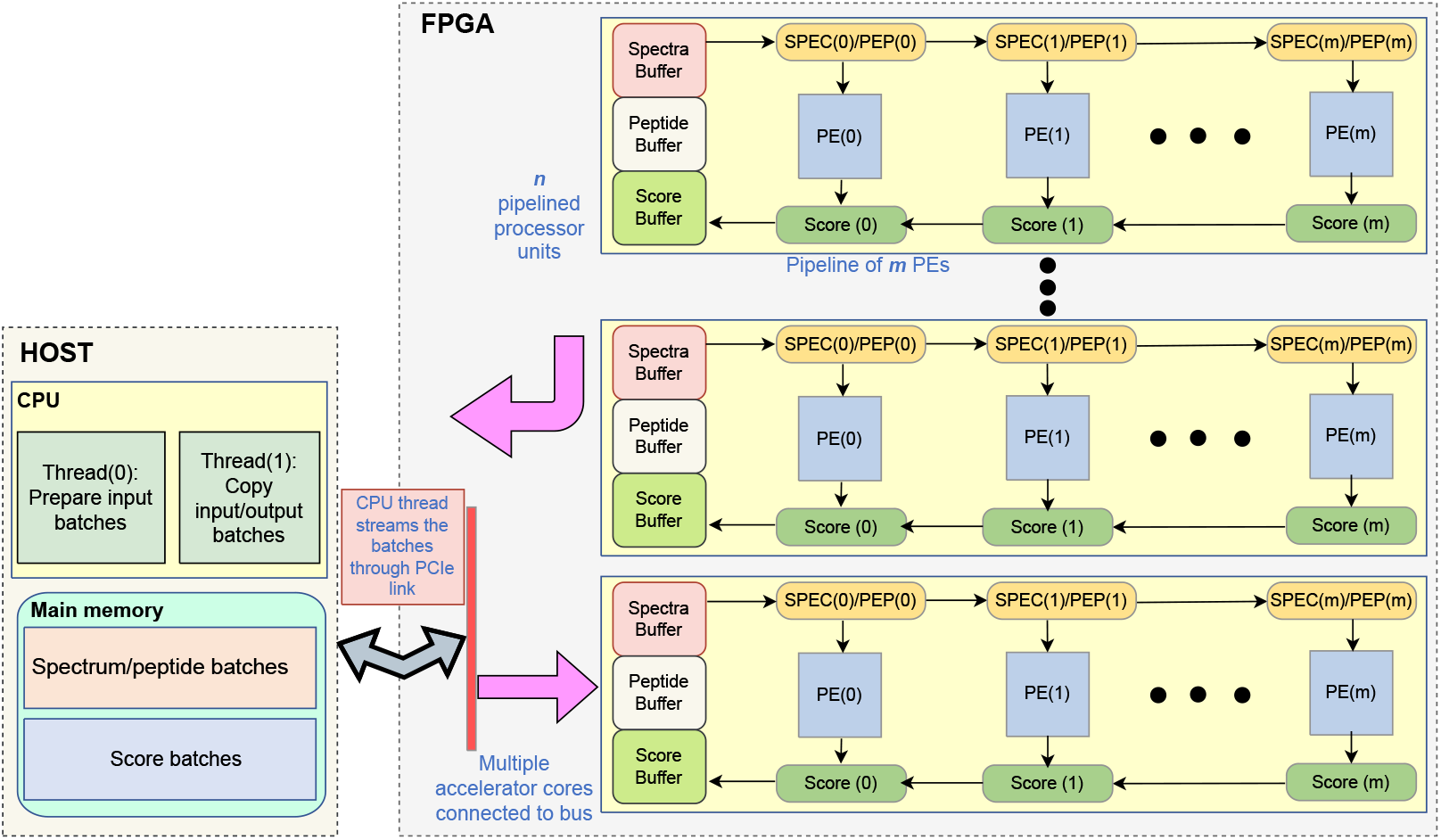
High-level organization of the accelerator system. Host CPU communicates with the FPGA via a PCIe link. All n pipelined processing units (PUs) are exposed to the CPU host. CPU periodically polls the PUs, and copies the respective spectra, peptides, or scores. Each PU is composed of a pipeline of m PEs through which peptides and scores are streamed.

### 2) Processing element to compute dot-score

To accelerate the dot-product computation in the inner-most loop, we designed and developed a configurable processing element shown in Fig. 4. Each processing element (PE) stores an experimental spectrum vector, and its corresponding candidate peptides in a fast on-chip RAM. To unroll the dotproduct computation in an inner-loop, each PE instantiates parallel dot-scorer modules. The number of dot-scorer modules is a selectable parameter that can be chosen at compile time and represents the loop-unrolling factor.

**Fig. 4.**
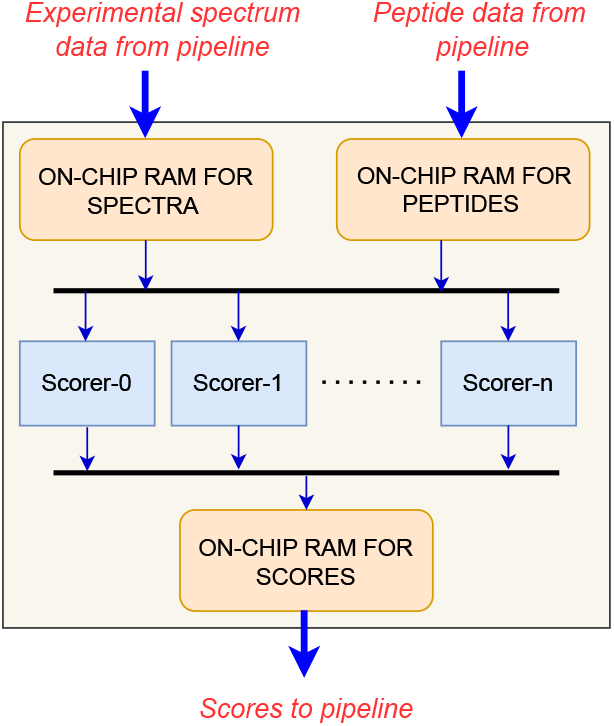
Architecture of the configurable processing element. An FSM controller reads the spectrum packets and peptide packets from the pipeline, and stores them in local on-chip RAMs. The scorer modules read the data from the bus and compute the dot-product concurrently, and the number of scorers is configurable according to the loop-unroll factor.

### 3) Design of dot-scorer module

#### Scorer Unit

The dot-scorer shown in Fig. 5 is made of two computational units namely theoretical-ion generator and ion matching unit. Theoretical ion generator computes the theoretical spectrum ions and an ion-matching unit matches the ions with experimental vectors, and generates scores. The ion-generator always has two peptides loaded into registers to implement double buffering which avoids computation stalls. On every clock cycle, the controller generates control signals for peptide sequence RAM to outputs the mass of two aminoacids – one corresponding to *N* terminus and other to the *C* terminus. The molecular masses from the *N* and *C* terminus are accumulated to generate the mass-to-charge ratio values of *b* and *y* ions which are then stored into the registers. This entire computation is performed by a combinational circuit enabling production of a new *b* or *y* ion in every clock cycle. The 2-to-1 mux in the design selects the theoretical ion which is then compared with the b- and y-ions of the experimental spectra in ascending order. All the new theoretical ions are simultaneously compared with all the experimental spectra ions utilizing parallel comparators in the ion-matching circuit.

**Fig. 5.**
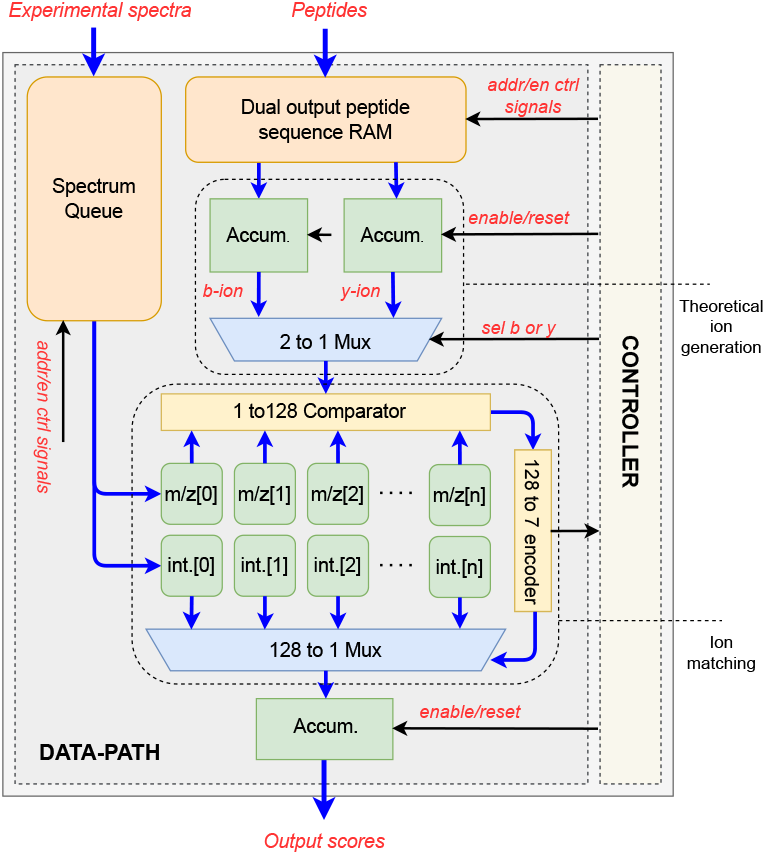
Scorer module design. Each theoretical ion is matched in parallel against all experimental ions using comparators, producing a bitmap of matches. The bitmap is encoded to select the matched ion via a multiplexer, and its intensity is accumulated into the peptide score.

#### Ion Matching Unit

During the score computation process, the ion-matching circuit is holding all the experimental spectrum ion pairs (m/z and intensity values) in registers. The m/z values of the theoretical ion are compared to the m/z values of the experimental ions. If a match occurs, the corresponding intensity value of that experimental ion is accumulated. To identify which ion from the experimental spectrum matched with the theoretical ion, a 128-to-7 encoder is used. Since the comparator will output a 1 when the match occurs, the encoder converts the 128 bitstream into a 7-bit value identifying the register index of the matching ion. For a given match, the encoder value will be greater than 1 which is then used to count the number of matches between the peptide and the experimental spectra. The accumulated intensity values along with the number of matched ions are bundled together to form the score packet which are stored in double-buffered registers. The score packets are then sent to the host machine to compute the final hyperscore.

### 4) Pipeline unit to process multiple spectra

The computation in the outer loops is limited by memory bandwidth. We therefore propose a pipelined processing unit (PU) shown in Fig. 3 that instantiates a pipeline of PEs to process multiple experimental spectra in parallel. This allows data reuse because mass-sorted experimental spectra share most of the candidate peptides. The PU generates three separate pipelines to move data to/from the PE pipeline. A spectrum pipeline, and a peptide pipeline to copy experimental spectra and candidate peptides to the PEs, and a score pipeline to read back the calculated scores. Each PU also has two on-chip RAM modules serving as private caches. One onchip RAM buffer holds the set of experimental spectra, and the other holds the candidate peptides along with computed scores. The PEs inside the accelerator core operate in a producer-consumer fashion i.e., collect experimental spectra and candidate peptides from the feeder pipelines and dispatch computed scores on the drain pipeline.

### C. Step 3: Design space exploration

Step 1 and 2 enables us to develop a highly parallel peptide deduction system. However, the performance of implemented design depends on several parameters such as nature of workload, architecture configuration, and dataflow scheme. Naturally, a subset of these architectural parameters will ensure maximal performance. To select the parameters that exhibit maximal performance, we need to enumerate all feasible design points for our given architecture. This design space exploration (DSE) over the architectural space which maximizes the performance is selected as the final design configuration.

To perform DSE quickly, we designed and developed an analytical performance model for the architecture by deriving latency equations for each design component. For a given set of workload and design parameters, the performance model estimates the total processing time, and compute/memory resource requirements of the FPGA. This enables us to define a constrained optimization problem, with total processing time as the objective function, and resource budget as constraints.

### 1) Parameter Space

The performance model is parameterized by both workload characteristics and architectural design choices. The workload is defined by the number of experimental spectra *M*, the number of candidate peptides per spectrum *k* (a function of the precursor mass window *δ*), and the peptide length *L*. The architectural configuration is defined by the number of processing units (PUs) *m*, processing elements (PEs) per PU *n*, loop unrolling factor *l*, and the spectrum and peptide batch sizes *S*_*sb*_ and *S*_*pb*_, respectively.

### 2) Performance Model

The execution latency of each architectural component is deterministic, enabling analytical estimation of the total execution time. As shown in Fig. 5, each PE implements a two-stage pipeline consisting of fragment-ion generation followed by dot-product computation. For b/y-ion searches, each amino acid generates two fragment ions, resulting in a per-peptide computation latency of 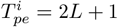 cycles.

All PEs within a PU operate concurrently on distinct candidate peptides. The time required for a peptide-compute operation (PCO) is therefore determined by the maximum PE latency across all *n* PEs and is given by

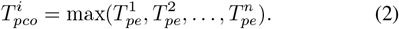

For each experimental spectrum, the PCO is repeated across all *k* candidate peptides. We define the spectrum-compute operation (SCO) as the sequence of PCOs required to process one spectrum. In addition to computation, peptide transfer operations incur communication overhead. Specifically, peptide-transfer (PCOM) and peptide-batch transfer (PBCOM) operations account for on-chip and off-chip data movement, respectively. Given a peptide batch size *S*_*pb*_, the time to process one spectrum is

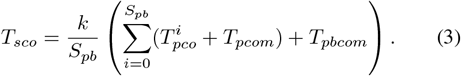

Each PU processes a subset of the *M* spectra. The time taken by the *j*-th PU to process the assigned spectra is obtained by summing the corresponding SCOs:

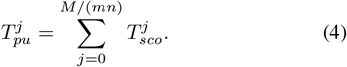

Similar to peptide transfers, spectrum processing incurs communication overhead through spectrum-transfer (SCOM) and spectrum-batch transfer (SBCOM) operations. For a spectrum batch size of *S*_*sb*_, the total processing time for the *j*-th PU is

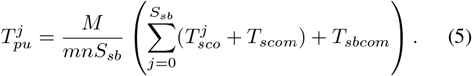

Since all PUs operate concurrently, the total execution time is determined by the slowest PU:

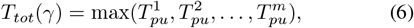

where *γ* = (*M, k, S*_*sb*_, *S*_*pb*_, *L, m, n*) denotes the parameter vector.

### 3) Resource consumption model

On-chip block memory is a constrained resource on FPGA and SoC platforms and directly limits exploitable parallelism. In FiCOPS, each processing element (PE) stores experimental spectra in local block RAM, while additional memory is consumed by spectrum and peptide buffers at the processing-unit (PU) level. We therefore restrict the design space to configurations that satisfy the available memory budget of the target FPGA.

Each PE stores two experimental spectra, each PU stores 2*S*_*sb*_ experimental spectra, and 4*S*_*pb*_ peptide and score entries. If *N*_*s*_ and *N*_*p*_ denote the number of bits required to store an experimental spectrum and a peptide/score, respectively, the total required on-chip memory for a configuration *γ* is estimated as

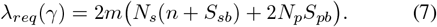

### D. External bandwidth model

Off-chip bandwidth requirements are determined by the frequency and size of CPU–FPGA data transfers. In the FiCOPS dataflow, three transfers dominate: (i) candidate peptide batches from CPU to FPGA, (ii) computed scores from FPGA to CPU, and (iii) experimental spectra from CPU to FPGA. For peptide batch processing time *T*_*pbc*_ and spectrum batch processing time *T*_*sbc*_, the required off-chip bandwidth is

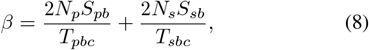

where 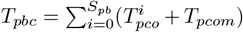 and 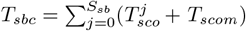.

### 1) Defining the optimization problem

The performance model enables formulation of the architectural design problem as a constrained optimization. Given workload parameters (*M, k*) and design parameters *γ* = (*S*_*pb*_, *S*_*sb*_, *L, m, n*), the objective is to minimize total execution time

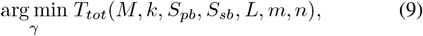

subject to FPGA resource constraints

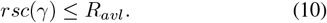

### 2) Design Parameters used in FICOPS

To find an optimal design, we explored the design space using the performance model and plotted a pareto frontier (see supplemental Fig. S1), where each data-point represents one design configuration. The pareto front represents a trade-off between resource utilization and execution time. By optimizing for various resource budgets, we found 8 different *design points*, and their implementation results are shown in table I. Next, we performed a parameter sweep across each design point for each variable while keeping the other variables constant, which resulted in implementation of 70 design configurations on the FPGA. The performance results are illustrated in Fig. 6 which show that increasing loop-unrolling and number of PUs increases the communication stall time, whereas increasing number of PEs steadily improves performance as it enables peptide reuse in the pipeline. For (*L, n*) *>* 3, there is no search time reduction as the decrease in computation time is complemented by an increase in communication stall time. Table I shows that the implementation size is limited by total adaptive logic modules (ALM) in the FPGA. The final design utilizing maximum resources instantiates 160 PEs, 3 PUs, and does not unroll the innermost loop.

**TABLE I.**
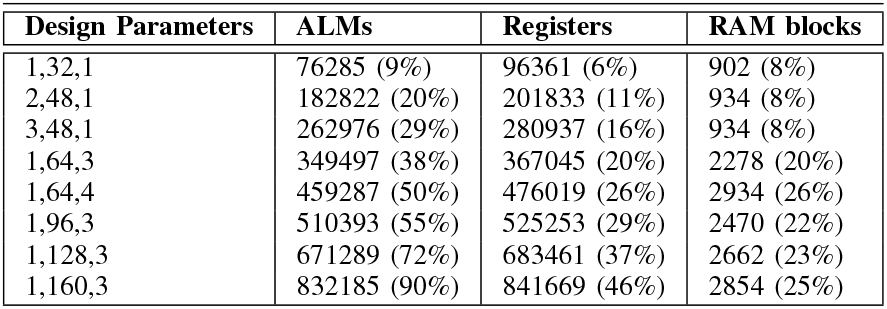
Design configurations identified for 10 different resource budget values.

**Fig. 6.**
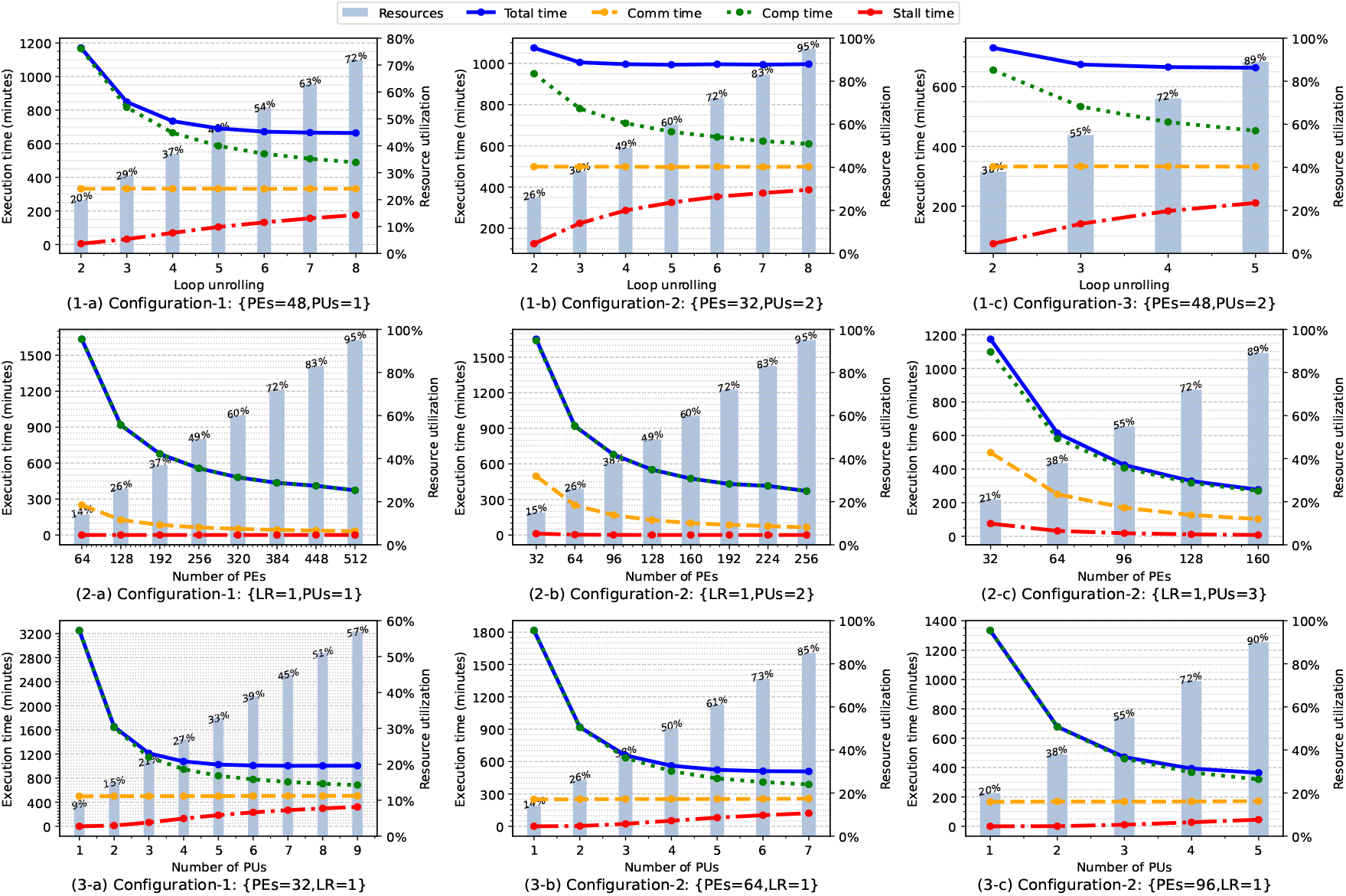
The effect of design parameters on total search time for PXD020590 dataset against human proteome database. (1) Increasing loop unrolling in dot scorer results in higher communication overhead due to added stress on memory bus. (2) Increasing PEs results in increases data reuse and reduces communication overhead. (3) Increasing PUs reduces search time when higher number of PEs are instantiated, but reduction plateaus when more than 3 PUs are instantiated due to increase in communication overhead.

Our DSE process reveals that using a simple PE to form a larger systolic array achieves better performance than designing a complex PE with more resources. To compare the pareto analysis results obtained from the performance model, we plotted the pareto front for execution time vs resources and execution time vs power consumption using the data of 70 designs implemented on FPGA (see supplemental Fig. S1). These curves demonstrate that the pareto curve from FPGA data closely follows the pattern of our performance model predictions. Furthermore, both kind of trade-off experiments (time vs resources and time vs power) would elect the same configuration as the most efficient design configuration. All of these results show that our performance modelling with resources, and parameters gives a good design metric that can be used for real implementation on FPGA and SoC boards.

## IV. Results

### A. Experimental Setup Overview

We compiled the FICOPS design configuration with 160 pipelined PEs and 3 processing units and synthesized the architecture for 200MHz clock speed. The design was implemented on an Intel Stratix 10 FPGA connected to the PCIe bus of the CPU system. The CPU used in this experiment was an octa-core Intel Core i7-4770 running at 3.40GHz with 16GB external memory.

#### 1) Datasets

For evaluation and comparison with existing solutions, we selected the following experimental spectra datasets from the ProteomeXchange database which comparable to the data sets used by HiCOPS [6] and GiCOPS [22] for evaluation. These data sets are PXD015890 (3.7M spectra), PXD020590 (1.6M spectra), PXD000612 (900K spectra), PXD009072 (300K spectra), PXD013332 (138K spectra), and PXD007871 (195K spectra).

#### 2) Database

All of the experiments were conducted using a protein sequence database for Homo sapiens from Uniport proteome ID UP000005640 (H. sapiens). We generated the peptides of length ranging from 6 to 50 with at most 2 missed cleavages by performing in-silico tryptic digestion of the protein sequences. For constructing the database, we selected the four most employed post-translational modifications (phosphorylation, oxidation, methylation and acetylation), which resulted in generation of 586 million peptides. The maximum modification per peptide is equal to 5.

### B. Search Time Comparison

Any database search engine can be broken-down into three distinct phases: (1) Database generation based on the search parameters (*database generation*); (2) processing of the spectra to make it capable of matching with the database entries (*spectra pre-processing*); (3) matching spectra to the database (*search*). Depending on what the search-engine is designed to accomplish, and how the processing is completed can vary between different tools with some tool performing better than others in some of the steps. To characterize the execution time for these steps, we performed the experiments and observed the times for each of the stages – both for closed and open searches – for single CPU tools (MSFragger [15], Crux [14], X!Tandem [23]), parallel CPU-only tools (HICOPS [6]) and CPU-Accelerator tools (FICOPS, GPU-Tide [24], GICOPS [22]). For serial experiments Crux was used for closed-search and MSFragger was used for open-search for fair comparisons. The total execution time for open search and closed search experiments against 6 benchmark datasets is plotted in Fig. 8, respectively. Each sub-figure shows a breakdown of total search-time which is divided into database construction, spectra pre-processing, and score computation. We discuss the breakdown of these run times.

**Fig. 7.**
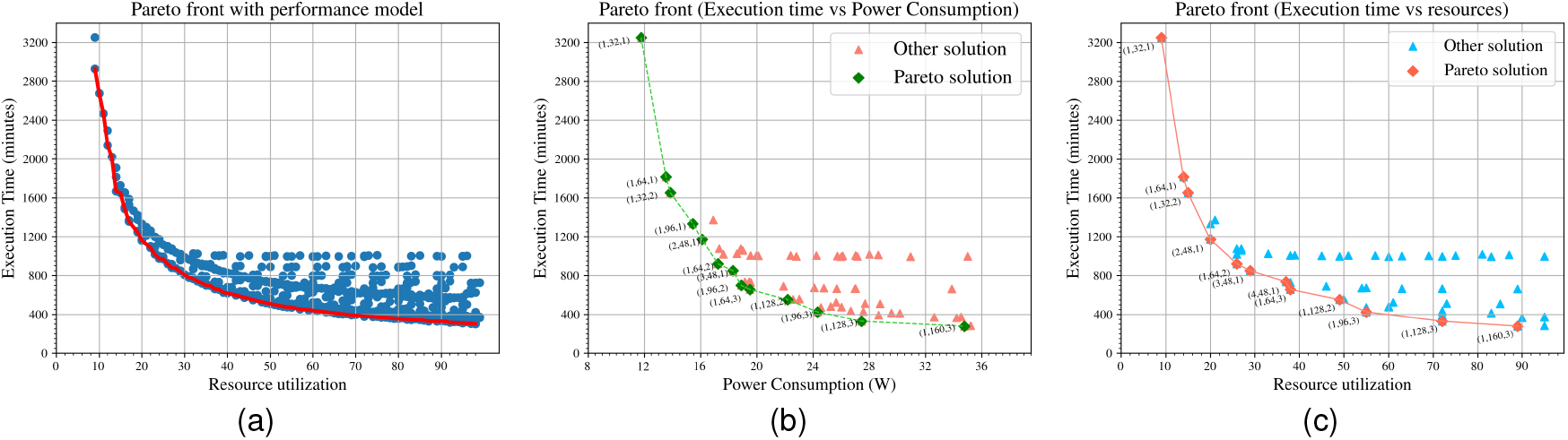
Pareto analysis reveals that increasing resource utilization linearly yields near linear speed-ups suggesting highscalability of design. (a) Performance model data (b) Time vs resources for 70 points (c) Time vs power for 70 points

**Fig. 8.**
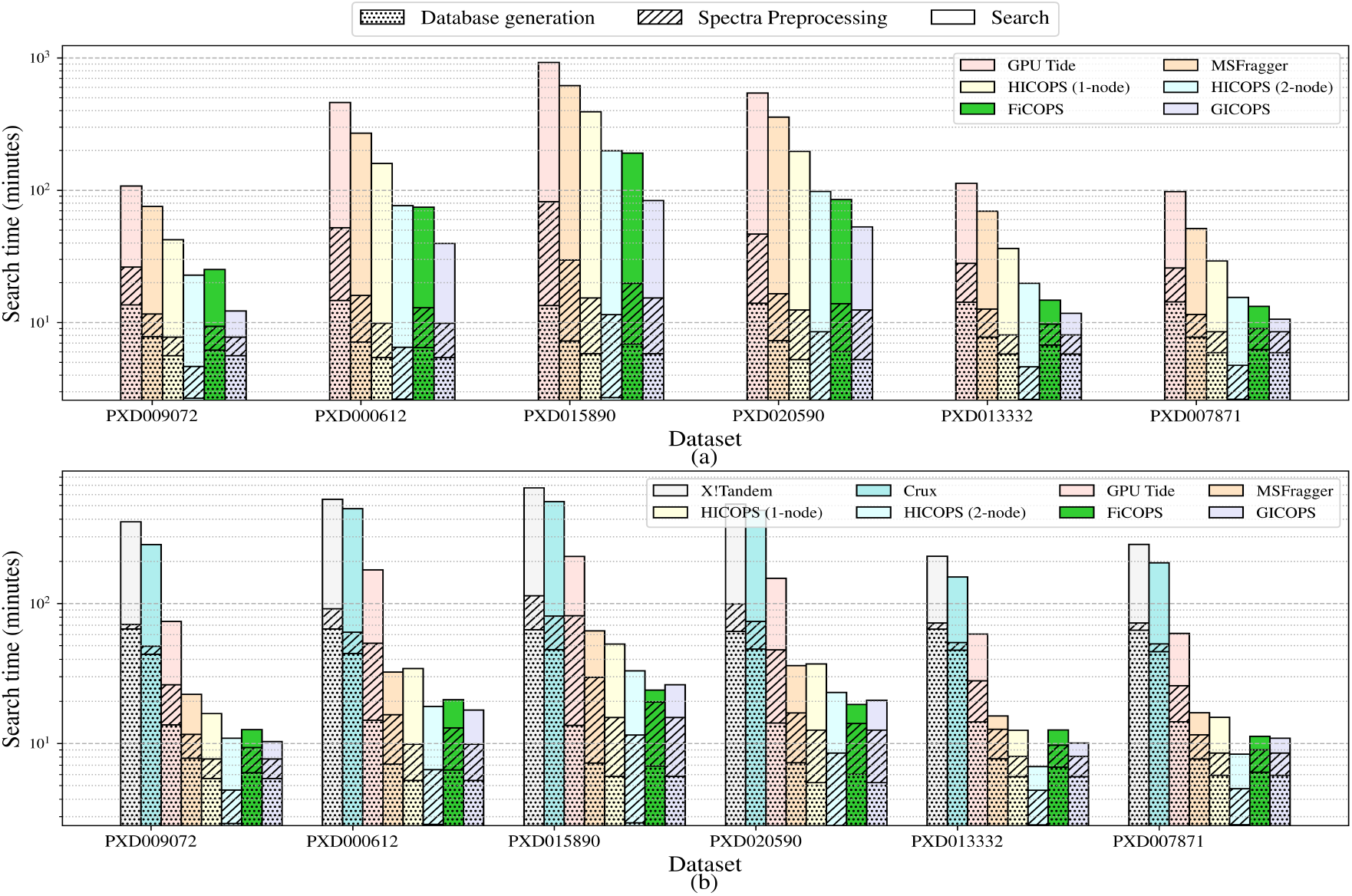
Comparison of run-time breakdown for database search against six benchmark proteomics datasets. In all the cases, FICOPs is faster than HICOPS (2-node) and MSFragger.(a) For open search experiments, time spent on search is atleast 5 times higher than combined time spent on pre-processing and database generation. (b) For smaller workloads in closed-search experiments, search time is atmost 3 times higher than pre-processing and database generation stages.

#### 1) Comparison with CPU-only Serial Tools

For open-search FICOPS is compared with MSFragger. <u>Database generation.</u> On average, FICOPS (6.42 min) performs better than MSFragger (7.50 min) for database generation. <u>Spectra Pre-processing.</u> On average, FICOPS (6.0 min) is able to outperform MSFragger (8.8 min) for spectra pre-processing. <u>Search.</u> On average, FICOPS (54.8 min) can outperform MSFragger (223.6 min) for searching the spectra against the database.

For closed-search FICOPS is compared with MSFragger, Crux and X!Tandem. <u>Database generation.</u> On average, FICOPS (6.42 min) runs much faster for database generation than X!Tandem (64.9 min), Crux (45.7 min), and comparable to MSFragger (7.50 min). <u>Spectra Pre-processing.</u> On average, FICOPS (6.02 min) is able to outperform all of the existing algorithms including X!Tandem (21.9 min), Crux (16.5 min), and MSFragger (8.8 min). <u>Search.</u> On average, FICOPS (4.2 min) was able to outperform X!Tandem (346.67 min), Crux (285.81 min), and MSFragger (14.8 min) by a wide margin.

#### 2) Comparisons with CPU-only Parallel Tools

For open-search FICOPS is compared with HICOPS. <u>Database generation.</u> On average, FICOPS (6.42 min) performs comparable to HICOPS 1-node (5.63 min) but subsequently HICOPS can perform better for 2-node (2.65 min) and 4-nodes (1.91 min). This is to be expected since HICOPS is a highly distributed framework, and the database generation can be accomplished independently on multiple nodes. <u>Spectra Pre-processing.</u> On average, FICOPS (6.0 min) performs comparably with HICOPS 1-node (4.7 min). However, HICOPS is not able to retain the advantage even with a greater number of nodes HiCOPS 2-node (4.1 min), and 4-node (3.83 min). <u>Search.</u> On average, FICOPS (54.8 min) outperforms HICOPS 1-node (132.2 min), and 2-node (65 min). HICOPS for 4-node (29.1 min) was able to outperform the FICOPS numbers as expected.

For closed-search FICOPS is compared with HICOPS. <u>Database generation.</u> On average, FICOPS (6.42 min) is comparable to HICOPS 1-node (5.63 min), but increasing the number of nodes results in better performance from HICOPS with node-2 (2.65 min) and 4-node (1.91 min), showcasing the scalability of the framework. <u>Spectra Pre-processing.</u> On average, FICOPS (6.02 min) is close to HICOPS 1-node (4.7 min), 2-node (4.11 min), and 4-node (3.83 min). While FICOPS seem to perform less than HICOPS, increasing the number of nodes for HICOPS does not dramatically improve the performance leading us to believe that FICOPS timing for many of the data sets will be comparable to HICOPS even when used in large supercomputing machines. <u>Search.</u> On average, FICOPS (4.2 min) can outperform HICOPS 1-node (17.5 min), 2-node (10.0 min), and 4-node (6.41 min). This is despite the fact that HICOPS is executed on advanced Intel processors for all nodes.

#### 3) Comparisons with CPU-Accelerator Parallel Tools

For open-search FICOPS is compared with GPU-Tide and GiCOPS. <u>Database generation.</u> On average, FICOPS (6.42 min) performs comparable to GICOPS (5.63 min) but outperforms GPU-Tide (14.04 min) by a large margin. <u>Spectra Pre-processing.</u> On average, FICOPS (6.01 min) performs comparably with GICOPS (4.7 min) but outperforms GPU-Tide (29.3min) by a large margin. <u>Search.</u> On average, FICOPS (54.8 min) performs a little less than GICOPS (24.7 min) but outperforms GPU-Tide (330.4 min) by a very large margin showcasing the scalability of our approach.

For closed-search FICOPS is compared with GPU-Tide and GiCOPS. <u>Database generation.</u> On average, FICOPS (6.42 min) is comparable to GICOPS (5.63 min) but outperforms GPU-Tide (14.1 min) showcasing the scalability of FICOPS approach despite low clock processing elements. <u>Spectra Pre-processing.</u> On average, FICOPS (6.02 min) performance is comparable to that of GICOPS (4.70 min) considering that CPU-GPU processing is accomplished using advance CPU and GPU architectures. Despite such an architecture FICOPS, which utilizes low-clock speeds, outperformance GPU-Tide (29.37 min) by a large margin. <u>Search.</u> On average, FICOPS (4.2 min) is comparable to GICOPS (5.53 min) but greatly outperforms GPU-Tide (79.52 min) by a wide margin.

Remarks. There are few observations that stand out for both closed-search and open-search experiments. One is that FiCOPS gives a better overall performance as compared to existing state of the art MSFragger (highly optimized serial method), and GPU-Tide which utilizes CPU-GPU architectures to process data. Furthermore, we can observe the overall performance of FICOPS in terms of execution time is close to GiCOPS which is a state-of-the-art method for processing MS data over memory-distributed clusters that are equipped with GPU’s. All of this is accomplished with smaller number of hardware and power resources that we demonstrate in the next sections.

### C. Speedup Comparisons with State of the Art

Once we were comfortable with the deduced peptides and the execution times, the next steps were to assess the speedups we will get from FICOPS, I.e., if the proposed hardware/software co-design is more scalable when compared to the existing state of the art methods. We selected MSFragger, HICOPS, GPU Tide, X!Tandem, and Crux for the comparisons since each of these tools provides a unique performance advantage. Both open-search and closed-search experiments were performed, depending on tool capabilities. The experiments, both for open and closed search, were categorized into CPU-only (serial), parallel (CPU-only), and parallel (CPU-GPU). Experiments using single node CPU (MSFragger, X!Tandem, Crux) were executed on Intel Xeon 6152 (22 cores) running at 2.10GHz, GPU-Tide was executed on Nvidia Titan Xp, and HICOPS (parallel CPU-only) was run on an in-house 6-node compute cluster built with Intel Xeon 4208 (16 cores each). Search experiments for all 6 datasets were executed using open-search setting (precursor window: 100Da), and closed-search (Precursor window: 50ppm) setting. Crux and X!Tandem were not included in the open-search experiments because each attempt resulted in the process to be killed after 24 hours of computations. Also both Crux and X!Tandem are not designed to perform open-search experiments.

#### 1) Comparison with CPU-only Serial Tools

FiCOPS was compared with X!Tandem, Crux and MSFragger as tools that employ whatever CPU is available to them. For closed search, FICOPS is 30.5×, 27×, 27.9×, 28×, 17× and 23× times faster as compared to X!Tandem. These are significant speedups with compared with other modern CPU tools such as Crux which exhibits on average 1.28×, and MSFragger which exhibits 14.8× speedups as compared to X!Tandem. Also note that these tools were executed on modern Xeon 6152, 2.10GHz CPUs with advance pipelining, Turbo Boost Max Technology, and Advance Vector Extensions. Even with these advanced processors, FICOPS was able to exhibit 2.99×, 3.62×, 3.23×, 4.2×, 4.7×, 3.88× speeds ups when compared with MSFragger for closed search operations.

#### 2) Comparisons with CPU-only Parallel Tools

For parallel CPU-only tools, HICOPS was selected as this was the only tool that offered a strategy that parallelizes both the database and the spectra. When comparing for closed search HICOPS as compared to X!Tandem exhibits, on average, 16.9×, 28.5×, and 37.2× speedup using 1, 2× and 4× nodes respectively. FICOPS exhibits 25.6× speedup when compared with X!Tandem. While this is at par with HICOPS (2 nodes), from Table II it can be seen that FICOPS exhibits speedups that are better than the 2-node HICOPS and comparable with HICOPS (with 4 nodes). Reduction in the average speedup in FICOPS is attributed to few of the datasets. The interesting part of these experiments were that FICOPS had similar characteristics for open-search results as well with 3.8× speedups, on average, when compared with MSFragger. For the same open-search experiments HICOPS exhibited 1.75× (1 node), 3.4× (2 node), and 6.8× (4 nodes) speedups. This shows that FICOPS consistently exhibits speedups that are better than the 2-node parallelization, and somewhat comparable with 4-node parallelization (depending on the data set) even when these tools are executed on advanced CPU architectures

**TABLE II.** Speedup performance of FICOPS over existing solutions. To ensure fair comparison X!Tandem search time is used to calculate the speedup for closed search experiments and MSFragger search time in used for open search experiments.

| Dataset | PXD009072 |  | PXD000612 |  | PXD015890 |  | PXD020590 |  | PXD013332 |  | PXD007871 |  |
| --- | --- | --- | --- | --- | --- | --- | --- | --- | --- | --- | --- | --- |
| Method | Open | Closed | Open | Closed | Open | Closed | Open | Closed | Open | Closed | Open | Closed |
| CPU-only Serial Tools |  |  |  |  |  |  |  |  |  |  |  |  |
| X!Tandem |  | 1 |  | 1 |  | 1 |  | 1 |  | 1 |  | 1 |
| Crux |  | 1.45 |  | 1.16 |  | 1.25 |  | 1.11 |  | 1.41 |  | 1.35 |
| MSFragger | 1 | 17.09 | 1 | 17.13 | 1 | 10.5 | 1 | 14.28 | 1 | 13.85 | 1 | 15.94 |
| CPU-only HPC Tools |  |  |  |  |  |  |  |  |  |  |  |  |
| HICOPS (1-node) | 1.78 | 23.45 | 1.69 | 16.14 | 1.58 | 13.03 | 1.81 | 13.87 | 1.92 | 17.54 | 1.76 | 17.14 |
| HICOPS (2-node) | 3.31 | 35.22 | 3.53 | 30.18 | 3.11 | 20.26 | 3.64 | 22.18 | 3.51 | 31.8 | 3.32 | 31.44 |
| HICOPS (4-node) | 7.91 | 38.99 | 6.52 | 39.07 | 7.06 | 30.97 | 6.94 | 33.25 | 7.26 | 37.98 | 5.25 | 42.99 |
| CPU-GPU and CPU-FPGA Tools |  |  |  |  |  |  |  |  |  |  |  |  |
| GiCOPS | 6.16 | 37.17 | 6.82 | 31.98 | 7.4 | 25.45 | 6.74 | 25.24 | 5.91 | 21.66 | 4.85 | 24.21 |
| GPU Tide | 0.7 | 5.15 | 0.59 | 3.2 | 0.67 | 3.08 | 0.66 | 3.4 | 0.62 | 3.6 | 0.53 | 4.31 |
| FiCOPS | 2.99 | 30.56 | 3.62 | 27.04 | 3.23 | 27.85 | 4.2 | 27.04 | 4.7 | 17.44 | 3.88 | 23.47 |

#### 3) Comparisons with CPU-Accelerator Parallel Tools

Finally, we completed our comparison with CPU-accelerator architectures. To this end, we choose GPU-Tide, and GiCOPS, both of which were run using CPU and a single GPU. While GICOPS is capable of exploiting parallelism on multiple

GPU’s on a single or multiple node, for these experiments only a single CPU-GPU was selected for fair comparison with FICOPS. For open-search GPU-Tide was able to perform with 0.63x speedup showcasing that it is slower to use GPU-Tide as compared to MSFragger. FICOPS exhibited 3.8× speedup and GICOPS speed up of 6.3× as compared with MSFragger. For closed search, FICOPS exhibited 25.6× speedup as compared to X!Tandem. This is comparable with GICOPS, a framework specifically designed for high-performance processing, a speedup of 27.6×. In contrast GPU-Tide exhibited 3.8× speedup as compared to X!Tandem on the same data sets – showcasing that using serial tools modified with a few CUDA directives do not scale well. In addition, it also shows that FICOPS is able to perform at the same pace and scalability as compared to a CPU-GPU architecture at a lower power footprint as we will discuss in the next section.

### D. Power Consumption Results

To perform power efficiency analysis, we measured the average power consumption of CPU solutions using the Intel RAPL interface. Power consumption for the GPU solution was measured using NVIDIA-smi during the execution, and an average over the period was calculated. For our FPGA design, we recorded the real time power usage values reported by the FPGA on-board power monitor interface.

Table III lists the average power consumption for all the solutions averaged across all 6 datasets for both closed search and open search setting. Moreover, for each experiment, we measured the total dot-product operations to find the throughput of each work in terms of giga operations per second (GOP/s) where the operation is a single dot-product computation. Using the power consumption and throughput, the power efficiency of in kilo operations per watt is also reported.

**TABLE III.** Power consumption, Throughput and Efficiency of both closed-and open-search experiments. For average power, lower is better, and for throughput and efficiency higher is more desirable.

| Method | Avg. Power (W) |  | Throughput (GOP/s) |  | Efficiency (kOPs/W) |  |
| --- | --- | --- | --- | --- | --- | --- |
|  | Open | Closed | Open | Closed | Open | Closed |
| **X!Tandem** | N/A | 109.258 | N/A | 0.012 | N/A | 0.1098 |
| **Crux** | N/A | 104.321 | N/A | 0.016 | N/A | 0.1533 |
| **MSFragger** | 106.77 | 115.365 | 2.14 | 0.651 | 20.0430 | 5.6429 |
| **HICOPS (1-node)** | 113.47 | 119.652 | 2.38 | 0.4925 | 20.9747 | 4.1161 |
| **HICOPS (2-node)** | 204.473 | 221.226 | 4.57 | 0.903 | 22.3501 | 4.0817 |
| **HICOPS (4-node)** | 199.546 | 411.647 | 8.28 | 1.484 | 41.4941 | 3.6050 |
| **GICOPS** | 176 | 196 | 10.99 | 1.391 | 62.4431 | 7.0969 |
| **GPU Tide** | 132 | 156 | 0.36 | 0.093 | 2.7272 | 0.5961 |
| **FiCOPS** | 32.769 | 34.537 | 7.49 | 0.77 | 228.57 | 22.29 |

#### 1) Comparison with CPU-only Serial Tools

For open-search FICOPS is compared with MSFragger. FICOPS (Avg Power: 32.8W) is able perform better than MSFragger (106.8 W) while outperforming on all computational segments (see Table III). This is also demonstrated when comparing FICOPS’s throughput (7.49 GOP/s) and Efficiency (228.6 KOP/W) as compared to MSFragger (2.14 GOP/s and 20.0 KOP/W). Similarly, closed search experiments also reveal, that FICOPS (34.5 W, 0.77 GOP/s, 22.3 KOP/W) outperforms MSFragger (115.4 W, 0.651 GOP/s, 5.6 kOPs/W), X!Tandem (109.3 W, 0.012 GOP/s, 0.1 kOPs/W), and Crux (104.32 W, 0.016 GOP/s, 0.153 kOPs/W) while exhibiting the same peptide deduction results in comparable wall clock time.

#### 2) Comparisons with CPU-only Parallel Tools

Both for open-search and closed-search FICOPS is compared with HICOPS. For open-search FICOPS (32.8 W, 7.49 GOP/s, 228.6 kOPs/W) is able perform better than HICOPS 1-node (113.47 W, 2.38 GOP/s, 20.97 kOPs/W), 2-node (204.5 W, 4.57 GOP/s, 22.35 kOPs/W), 4-node(199.5 W, 8.28 GOP/s, 41.49 kOPs/W) by a wide margin especially when considering that the wall-clock time achieved by HICOPS for all nodes is comparable to FICOPS. Similarly, closed search experiments also reveal, that FICOPS (34.5 W, 0.77 GOP/s, 22.3 KOP/W) outperforms HICOPS 1-node (119.7 W, 0.493 GOP/s, 4.12 kOPs/W), HICOPS 2-node (221.2 W, 0.903 GOP/s, 4.08 kOPs/W), and HICOPS 4-node (411.6 W, 1.48 GOP/s, 3.6 kOPs/W). These experiments conclusively demonstrate that FICOPS is more energy-efficient when compared to leading CPU-only HPC frameworks, while comparably scalable.

#### 3) Comparisons with CPU-Accelerator Parallel Tools

Both for open-search and closed-search FICOPS is compared with GPU-Tide and GiCOPS. For open-search FICOPS (32.8 W, 7.49 GOP/s, 228.6 kOPs/W) is able outperform in the average power as compared to GICOPS (176 W, 10.99 GOP/s, 62.44 kOPs/W). While GICOPS can outperform GOP/s, the efficiency (kOPs/W) is still less than that of FICOPS. In other words, FICOPS still outperforms in terms of the operations/wattage for a given experiment. FICOPS can outperform and GPU-Tide (132 W, 0.36 GOP/s, 2.73 kOPs/W) across the board for average power, throughput as well as efficiency. For closed search, FICOPS (32.8 W, 7.49 GOP/s, 228.6 kOPs/W) can outperform both GiCOPS (196 W, 1.391 GOP/s, 7.09 kOPs/W) and GPU-Tide (156 W, 0.093 GOP/s, 0.596 kOPs/W) in terms total power, giga operations per second as well as kilo operations per watt.

Remarks: For both search windows, FiCOPS consumes 3× less power than all CPU solutions, and 5× less power than GPU solutions. Further, FiCOPS achieves 10× higher performance/watt ratio than single-node CPU solutions, 5× higher than HICOPS running on 4 CPU nodes, and more than 100× higher than existing GPU solution, while exhibiting comparable wall clock times for searching – showcasing the scalability of FICOPS while being extremely energy efficient.

## V. Discussion

In this paper, we designed and developed a CPU-FPGA heterogeneous accelerator system for end-to-end peptide database search. To accomplish this task, we systematically analyzed the computational requirements, bottlenecks, and opportunities of parallelism in the workflow. Our in-depth analysis found that to efficiently accelerate database search tasks, without using the memory intensive fragment-ion index, redundant comparison operations in the inner-join process need to be executed in parallel, and the associated memory latency needs to be minimized. To this end, we developed a highly pipelined architecture that enabled minimization of communication overhead by overlapping computation and communication. We then redesigned the algorithm, and proposed a highly parameterized and novel architecture template to implement the score computation workflow.

The current serial tools struggle when confronted with large databases, development of HPC methods have been ignored or downright opposed by the mass spectrometry community – despite overwhelming evidence to the contrary. While serial algorithms such as Crux were optimized for old single-core systems, they have been left behind either due to lack of scalability on modern multicore and GPU architectures, or with the introduction of much more powerful open-search methods such as MSFragger. Part of the reason is increased interest in analysis of more complex data sets from human microbiome or environmental samples. Increased interest in detecting and searching of novel and uncommon PTM’s that may be involved in human health are also stretching the capabilities of current serial algorithms. Despite these limitations, these tools are useful for small data sets, and databases with limited PTM’s or limited number of species need to be investigated.

More recently, we have designed and developed highperformance computing tools, HiCOPS, and GiCOP that can accelerate and enable tera-scale database searching and computing using memory-distributed supercomputers, and GPU-enabled architectures, respectively. While steps in the right direction these frameworks used existing general-purpose CPU and GPU hardware to exploit parallelism and pipelining to accomplish the computations. To date, custom-designed hardware with FPGA’s that can process MS database search has been limited. For MS data and other domains, isolated section of a large workflow is often seen for hardware acceleration. However, integrating such hardware designs with rest of the software workflow stack is often challenging and has not been investigated by us or others. Again, part of the reasons on why this has not been accomplished in this MS data domain is due to large data sets that are required to be parsed and indexed and a naïve integration with the rest of the workflow diminishes any speedups gained by hardware accelerators and may even slow it down due to large volume of data. Although FPGA based accelerators are essential for software/hardware designs that can be incorporated into the mass spectrometry machines for processing of data – such investigations have been limited due to significant skillset (hardware/software co-design, FPGA, parallel processing, and MS/omics) and resources demanded by such development projects. Further the problem of designing hardware, and then programming it using low-level Verilog and VHDL is beyond the scope for many software engineering students and researchers. While OpenCL High-Level Synthesis (HLS) has been proposed and available, their performance as compared to the Hardware Descriptive Languages (HDL) is still subpar at best [23], [24].

In this study, we designed and developed a highly optimized hardware solution to the MS database search problem. The rationale for this FPGA based codesign is to develop customized hardware that can process the complex MS based omics workflows. To search for an optimal design that would work as a general-purpose hardware-based workflow, hundreds of designs needed to be evaluated which can be a time consuming and error prone process since compilation of a single complex FPGA design can take multiple hours. To circumvent this limitation, we developed an analytical performance model, and performed rapid design space exploration to find the trade-offs between area, and performance. Our design space exploration process revealed that generating a simpler, and smaller PE enabled instantiation of deeper pipelines. Since a complex PE puts more pressure on the memory bus, the performance gain achieved by loop unrolling was quickly diminished by increasing communication costs. Using a deeper pipeline of simple processing elements not only increased processing density but also enabled data reuse. The pareto analysis of our design showed linear scalability with with the number of resources available in the FPGA.

The implementation of this design and subsequent performance evaluation was accomplished for datasets with different fragmentations, species and databases. Our extensive experimentation demonstrated that FiCOPS achieves, on average, 3.5× speed-up against existing CPU solutions (HiCOPS [11]) while consuming 5.7× less power compared to to existing state-of-art GPU solutions (GiCOPS [22]). FiCOPS also exhibited superior watt ratio performance as compared to all existing serial or parallel solutions. From these results, we conclude that FPGA-based heterogeneous computing solutions offer a promising avenue for intensive mass spectrometrybased omics problems. Such frameworks will be especially useful for energy-efficient training and inference of new cohort of machine-learning models that are being developed and deployed (Specollate [25], ProteoRift [26]) for MS based omics.

This study and work can be used to demonstrate two main conclusions. One, that despite FICOPS using a lowspeed clocks and subsequently lower power, the proposed framework can compete with much larger GPU based GICOPS framework. In other words, it is not the hardware that is at your disposal that determines the speedup but how the computations are done for a given hardware, or in FICOPS case how the hardware is modelled to do the computations. Another thing this study teaches is that just throwing hardware (like GPU’s) at codebases like Tide/MSFragger to make it “GPU ready” does not necessarily equate to performance gains. On the contrary we can see that GPU-Tide is even slower than its serial (Tide/Crux) code base when used with GPU’s. This is likely because GPU-Tide does not consider communication and I/O costs associated with CPU-Accelerator architectures, which we have demonstrated in Fig. 8, are a major source of costs associated with peptide deduction search-engines. Some of the limitation of FICOPS includes its current focus solely on the search computation stage, while database generation and postprocessing are still performed on the CPU. However, these limitations can be addressed through continued development to achieve end-to-end processing workflow. Future efforts can develop hardware accelerators for scoring as well as the data wrangling part of the workflows that can then be integrated with the rest of the software stack that can be used in a laboratory setting.

## VI. Biography Section

**Sumesh Kumar** is a PhD student at the Knight Foundation School of Computing and Information Sciences at FIU, Miami FL USA. His research interests include hardware software co-design for computational biology and compute intensive ML problems.

**Joseph A. Zambreno** has been with the Department of Electrical and Computer Engineering at Iowa State University since 2006, where he currently holds the Ross Martin Mehl and Marylyne Munas Mehl Professorship in Computer Engineering. His research interests include computer architecture, reconfigurable computing, and hardware security.

**Ashfaq Khokhar** is a Professor and Palmer Department Chair of Electrical and Computer Engineering at Iowa State University. He received the Ph.D. degree in computer engineering from the University of Southern California in 1993. Khokhar’s research centers on health care data mining, contentbased multimedia modeling, retrieval and multimedia communication, highperformance algorithms, and context-aware wireless sensor networks.

**Shoaib Akram** is a Lecturer in the School of Computing at the Australian National University. He received the Ph.D. degree in Computer Science and Engineering from Ghent University. His research interests include computer architecture, hardware–software interfaces, and data-centric system optimization.

**Fahad Saeed** is a Professor in the Knight Foundation School of Computing and Information Sciences at Florida International University. He received the Ph.D. degree in Electrical and Computer Engineering from the University of Illinois at Chicago in 2010. His research interests lie at the intersection of machine learning, high-performance computing, and computational biology.

## Supplemental Information

### Related Work

This section summarizes prior efforts on accelerating peptide database search using parallel computing platforms. The reviewed works are grouped by the underlying hardware architecture, including multicore CPUs, distributed-memory systems, and graphics processing units (GPUs), and provide context for positioning the proposed FPGA-based approach relative to existing solutions.

### A. Acceleration with distributed computing

SEQUEST-PVM [1] explored distributed computing approach and implemented a parallel version of SEQUEST [2] using parallel virtual machine (PVM) framework to distribute the experimental spectra, and protein database files among nodes in the system to achieve near linear speed-ups (4.98× for 5 nodes). Another work (Parallel Tandem [3]) implemented the parallel version of X!tandem by following the same approach and used PVM and MPI protocols to distribute the search among 40× compute nodes to obtain 36-fold speed-up for only dot-product operation. However, the approach used by Parallel Tandem suffered from load imbalance which was improved by X!!Tandem using random shuffling and ownercomputes policy to achieve 29× speed-up with 64× nodes for the entire search application. MR-Tandem also parallelized X!Tandem and mimicked the approach used in X!!Tandem but used Hadoop instead of MPI to achieve fault-tolerance in their network. However, their approach was 20% slower than the X!!Tandem’s MPI based implementation. Moreover, Hydra [4] also used Hadoop and did a complete re-write of the X!Tandem framework to fully utilize the MapReduce architecture. All of these methods were based on embarrassingly parallel methodology which resulted in sub-optimal usage of disk-space and memory on each nodes. Therefore, these parallel strategies have limited performance improvement for larger databases for meta-proteomics and proteogenomics analysis for a given architecture. More recently, we have presented a HPC framework for acceleration of database search on distributed-memory supercomputers. The HiCOPS framework uses a novel approach in which the (massive) theoretical databases are partially distributed across parallel nodes in a load-balanced fashion followed by asynchronous parallel execution of the database peptide search on the partial database. On completion, the locally computed results are merged into global results in a communication-optimal manner. Since HiCOPS is introduced as a framework that is search-algorithm oblivious - the design can be extended or replaced to accelerate most existing and future search algorithms. HiCOPS is shown to perform tera-scale database computations with at least 10× speedup as compared to existing parallel solutions that exploit embarrassingly parallel methods.

### B. Acceleration with GPU

Given that Graphical Processing Units (GPU) are ubiquitous, its application in the database-search engine has been limited. FastPASS [5] used a Nvidia Tesla C1060 to implement spectral library search to achieve a 274× speed-up against X!Tandem. However, the work was limited by searching for only spectral library which is a much smaller database and does not incorporate PTMs. In Tempest, a CPU-GPU hybrid approach was proposed to implement cross-correlation function (Xcorr) – used in SEQUEST and Crux – to achieve 15× speed-up compared to a CPU only approach. The hybrid design of Tempest used CPU to perform protein digestion and offload candidate peptides to GPU which were then scored against a single experimental spectrum in parallel. This approach required multiple copies of the experimental spectrum to be made which introduced extra communication overhead. In [6], the spectrum dot-product between experimental and theoretical spectrum as a matrix-vector multiplication problem was introduced and offloaded it to a GPU. This design reported 30 to 60 times speed using a single GPU and demonstrated an almost linear scalability of their design on a GPU cluster. Another approach proposed in [7], distributed the fragment ions in the theoretical spectrum among GPU threads to perform a fine-grained computation to dot-product and achieved 2.7 times speed-up compared to Crux. While these techniques exploited parallelism for CPU-GPU architectures, similar to other parallel techniques, they have been limited by using embarrassingly parallel models, and limited speedups.

More recently, we developed GiCOPS, a CPU–GPU framework that extended the HiCOPS partitioned-search approach using techniques such as Savitzky–Golay smoothing and standard tail-fit models. Although GiCOPS achieved strong speedups on large CPU–GPU systems, its design relied heavily on HPC-scale memory, bandwidth, and coarse parallelism. These constraints limited its portability and made it less suitable for lighter or emerging architectures. These limitations motivated the development of FiCOPS, which targets finergrained parallelism and lower resource requirements.

### C. Acceleration with FPGA

While non-scalability is widely acknowledged, usage of custom-designed reconfigurable accelerators has not been extensively studied for peptides deduction and database search. Wang et al. [8] developed an accelerator architecture that instantiated six scoring modules, and one fragment ion generation module to obtain 174× speed-up compared to X!Tandem. The same group extended their design [9] as a CPU-FPGA co-design accelerator which enabled protein digestion calculations on the FPGA. While they presented their results using 2-FPGAs where the CPU was managing task distribution, only 10x speedups as compared to X!Tandem were realized. Further, no comparison with existing state-of-the-art serial methods (Crux, MSFragger) were presented and further investigation is warranted. More recently, our group implemented a data reuse strategy to reduce communication cost and achieved 24-fold speed-up compared to Crux which is reported as a preliminary work [10].

**TABLE S1.**
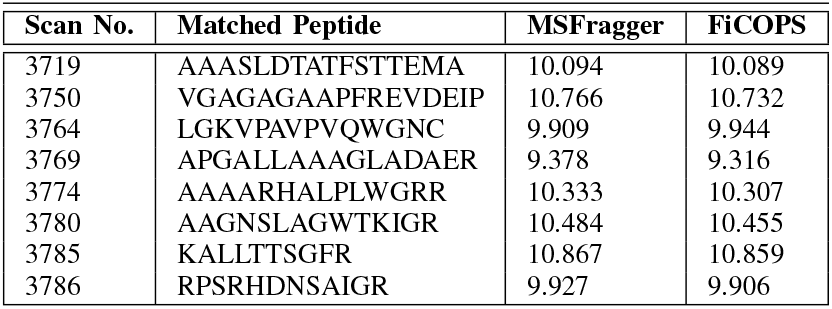
Comparison of computed hyperscore value between MSFragger and FiCOPS.

Remarks: Software/hardware co-design that can process the data in a scalable manner and in almost real-time is missing from the literature. Despite their immense potential, relatively few papers have been published using Field programmable gate arrays (FPGA) [8]–[10], potentially due to the complexity of designing and developing hardware/software co-designs for complex MS data. The proposed next generation of parallel computing algorithm development based on communicationavoidance, I/O reduction, decomposition, hardware exploits, and hardware/software design (for FPGA’s) will fill a critical gap in computational infrastructure that may lead to understanding and ability to deduce novel peptides and will contribute a fundamental tool for studying complex communities in proteomics, and meta-proteomics data.

## Correctness Analysis

The first assessment that we made was to ensure that the results that we are getting on FiCOPS is comparable to what one would get when using a serial algorithm such as MSFragger. While computing results and scores for distributed scoring mechanism, such is as done in HiCOPS, may result in some deviation from the serialized-score, we expect FiCOPS score to be within the error-bound of the hardware on which they were processed. We verified the consistency of results across parallel runs by searching the database against all the PXD data sets using various settings and PTM combinations. The correctness was evaluated in terms of identified peptide sequences, and the corresponding hyperscores and FiCOPS scores which were consistent within ±0.1. For all our experiments we found that the scores calculated using FiCOPS architecture were within a reasonable margin of error up to 3 decimal places which can be attributed to small number of bits used for FiCOPS architecture as compared to 64 bit machine used for MSFragger calculations. In addition, we demonstrate that the associated peptide match is the same for all the scores that have been calculated. A snapshot of the results is shown in Table S1.

The Table conclusively shows that the difference in the scores that are calculated using FiCOPS and the one calculated using MSFragger are minimal and not statistically meaningful, and would result in same peptide matches. This result is to be expected since FICOPS is designed to replicate the hyperscores calculations without any changes, or approximations.

## Notes

### Competing Interest Statement

The authors have declared no competing interest.

### Funding

- National Institutes of Health: R35GM153434
- U.S. National Science Foundation: OAC-2312599

## References

[1] M. Y. Ang, T. Y. Low, P. Y. Lee, W. F. W. M. Nazarie, V. Guryev, and R. Jamal, “Proteogenomics: from next-generation sequencing (NGS) and mass spectrometry-based proteomics to precision medicine,” Clinica chimica acta, vol. 498, pp. 38–46, 2019, publisher: Elsevier.

[2] M. U. Tariq, M. Haseeb, M. Aledhari, R. Razzak, R. M. Parizi, and F. Saeed, “Methods for proteogenomics data analysis, challenges, and scalability bottlenecks: a survey,” IEEE Access, vol. 9, pp. 5497–5516, 2020, publisher: IEEE.

[3] M. F. Savitski and M. M. Savitski, “Unbiased detection of posttranslational modifications using mass spectrometry,” Methods in Molecular Biology (Clifton, N.J.), vol. 673, pp. 203–210, 2010.

[4] M.-S. Kim, J. Zhong, and A. Pandey, “Common errors in mass spectrometry-based analysis of post-translational modifications,” PROTEOMICS, vol. 16, no. 5, pp. 700–714, 2016, eprint: https://onlinelibrary.wiley.com/doi/pdf/10.1002/pmic.201500355. [Online]. Available: https://onlinelibrary.wiley.com/doi/abs/10.1002/pmic.201500355

[5] J. K. Eng, B. C. Searle, K. R. Clauser, and D. L. Tabb, “A face in the crowd: recognizing peptides through database search,” Molecular & Cellular Proteomics, vol. 10, no. 11, 2011, publisher: ASBMB.

[6] M. Haseeb and F. Saeed, “HiCOPS: High Performance Computing Framework for Tera-Scale Database Search of Mass Spectrometry based Omics Data,” arXiv preprint arXiv:2102.02286, 2021.

[7] H. Li, Y. S. Joh, H. Kim, E. Paek, S.-W. Lee, and K.-B. Hwang, “Evaluating the effect of database inflation in proteogenomic search on sensitive and reliable peptide identification,” BMC Genomics, vol. 17, no. 13, p. 1031, Dec. 2016. [Online]. Available: 10.1186/s12864-016-3327-5

[8] K.-F. Wang, Y.-Z. Wu, and H. Chi, “A universal database reduction method based on the sequence tag strategy to facilitate large-scale database search in proteomics,” International Journal of Mass Spectrometry, vol. 483, p. 116966, Jan. 2023. [Online]. Available: https://www.sciencedirect.com/science/article/pii/S1387380622001713

[9] E. Boschetti and P. G. Righetti, “Low-Abundance Protein Enrichment for Medical Applications: The Involvement of Combinatorial Peptide Library Technique,” International Journal of Molecular Sciences, vol. 24, no. 12, p. 10329, Jan. 2023, number: 12 Publisher: Multidisciplinary Digital Publishing Institute. [Online]. Available: https://www.mdpi.com/1422-0067/24/12/10329

[10] K. H. Kim, Y. H. Ahn, E. S. Ji, J. Y. Lee, J. Y. Kim, H. J. An, and J. S. Yoo, “Quantitative analysis of low-abundance serological proteins with peptide affinity-based enrichment and pseudo-multiple reaction monitoring by hybrid quadrupole time-of-flight mass spectrometry,” Analytica Chimica Acta, vol. 882, pp. 38–48, Jul. 2015.

[11] L. E. Kilpatrick and E. L. Kilpatrick, “Optimizing High-Resolution Mass Spectrometry for the Identification of Low-Abundance Post-Translational Modifications of Intact Proteins,” Journal of proteome research, vol. 16, no. 9, pp. 3255–3265, Sep. 2017. [Online]. Available: https://www.ncbi.nlm.nih.gov/pmc/articles/PMC7489340/

[12] A. I. Nesvizhskii, “Proteogenomics: concepts, applications and computational strategies,” Nature methods, vol. 11, no. 11, pp. 1114–1125, 2014, publisher: Nature Publishing Group.

[13] R. Craig and R. C. Beavis, “Tandem: matching proteins with tandem mass spectra,” Bioinformatics, vol. 20, no. 9, pp. 1466–1467, 2004.

[14] S. McIlwain, K. Tamura, A. Kertesz-Farkas, C. E. Grant, B. Diament, B. Frewen, J. J. Howbert, M. R. Hoopmann, L. Käll, J. K. Eng, and others, “Crux: rapid open source protein tandem mass spectrometry analysis,” Journal of proteome research, vol. 13, no. 10, pp. 4488–4491, 2014, publisher: ACS Publications.

[15] A. T. Kong, F. V. Leprevost, D. M. Avtonomov, D. Mellacheruvu, and A. I. Nesvizhskii, “MSFragger: ultrafast and comprehensive peptide identification in mass spectrometry–based proteomics,” Nature methods, vol. 14, no. 5, pp. 513–520, 2017, publisher: Nature Publishing Group.

[16] H. Kim, S. Han, J.-H. Um, and K. Park, “Accelerating a cross-correlation score function to search modifications using a single GPU,” BMC Bioinformatics, vol. 19, no. 1, p. 480, Dec. 2018. [Online]. Available: 10.1186/s12859-018-2559-6

[17] A. I. Nesvizhskii, “A survey of computational methods and error rate estimation procedures for peptide and protein identification in shotgun proteomics,” Journal of Proteomics, vol. 73, no. 11, pp. 2092–2123, Oct. 2010. [Online]. Available: https://www.sciencedirect.com/science/article/pii/S1874391910002496

[18] G. C. Teo, D. A. Polasky, F. Yu, and A. I. Nesvizhskii, “A fast deisotoping algorithm and its implementation in the MSFragger search engine,” Journal of proteome research, vol. 20, no. 1, pp. 498–505, Jan. 2021. [Online]. Available: https://www.ncbi.nlm.nih.gov/pmc/articles/PMC8864561/

[19] D. J. Geiszler, D. A. Polasky, F. Yu, and A. I. Nesvizhskii, “Detecting diagnostic features in MS/MS spectra of post-translationally modified peptides,” Nature Communications, vol. 14, no. 1, p. 4132, Jul. 2023, publisher: Nature Publishing Group. [Online]. Available: https://www.nature.com/articles/s41467-023-39828-0

[20] J. K. Eng, B. Fischer, J. Grossmann, and M. J. MacCoss, “A Fast SEQUEST Cross Correlation Algorithm,” Journal of Proteome Research, vol. 7, no. 10, pp. 4598–4602, Oct. 2008, publisher: American Chemical Society. [Online]. Available: 10.1021/pr800420s

[21] J. K. Eng, T. A. Jahan, and M. R. Hoopmann, “Comet: an open-source MS/MS sequence database search tool,” Proteomics, vol. 13, no. 1, pp. 22–24, 2013, publisher: Wiley Online Library.

[22] M. Haseeb and F. Saeed, “GPU-acceleration of the distributed-memory database peptide search of mass spectrometry data,” Scientific Reports, vol. 13, no. 1, p. 18713, Oct. 2023.

[23] R. Craig and R. C. Beavis, “A method for reducing the time required to match protein sequences with tandem mass spectra,” Rapid communications in mass spectrometry, vol. 17, no. 20, pp. 2310–2316, 2003, publisher: Wiley Online Library.

[24] H. Kim, S. Han, J.-H. Um, and K. Park, “Accelerating a cross-correlation score function to search modifications using a single GPU,” BMC bioinformatics, vol. 19, no. 1, pp. 1–5, 2018, publisher: BioMed Central.

## References

[1] R. G. Sadygov, J. Eng, E. Durr, A. Saraf, H. McDonald, M. J. MacCoss, and J. R. Yates, “Code developments to improve the efficiency of automated MS/MS spectra interpretation,” Journal of proteome research, vol. 1, no. 3, pp. 211–215, 2002, publisher: ACS Publications.

[2] J. K. Eng, A. L. McCormack, and J. R. Yates, “An approach to correlate tandem mass spectral data of peptides with amino acid sequences in a protein database,” Journal of the american society for mass spectrometry, vol. 5, no. 11, pp. 976–989, 1994, publisher: ACS Publications.

[3] D. T. Duncan, R. Craig, and A. J. Link, “Parallel tandem: a program for parallel processing of tandem mass spectra using PVM or MPI and X! Tandem,” Journal of proteome research, vol. 4, no. 5, pp. 1842–1847, 2005, publisher: ACS Publications.

[4] S. Lewis, A. Csordas, S. Killcoyne, H. Hermjakob, M. R. Hoopmann, R. L. Moritz, E. W. Deutsch, and J. Boyle, “Hydra: a scalable proteomic search engine which utilizes the Hadoop distributed computing framework,” BMC bioinformatics, vol. 13, no. 1, pp. 1–6, 2012, publisher: BioMed Central.

[5] L. A. Baumgardner, A. K. Shanmugam, H. Lam, J. K. Eng, and D. B. Martin, “Fast parallel tandem mass spectral library searching using GPU hardware acceleration,” Journal of proteome research, vol. 10, no. 6, pp. 2882–2888, 2011, publisher: ACS Publications.

[6] Y. Li and X. Chu, “Speeding up scoring module of mass spectrometry based protein identification by GPU,” in 2012 IEEE 14th International Conference on High Performance Computing and Communication & 2012 IEEE 9th International Conference on Embedded Software and Systems. IEEE, 2012, pp. 1315–1320.

[7] Y. Li, H. Chi, L. Xia, and X. Chu, “Accelerating the scoring module of mass spectrometry-based peptide identification using GPUs,” BMC bioinformatics, vol. 15, no. 1, pp. 1–11, 2014, publisher: BioMed Central.

[8] J. Qiu, P. Kang, L. Ding, Y. Yuan, W. Yin, and L. Wang, “Fpga acceleration of the scoring process of x! tandem for protein identification,” in 2017 27th International Conference on Field Programmable Logic and Applications (FPL). IEEE, 2017, pp. 1–4.

[9] M. Yang, T. Chen, X. Zhou, L. Zhao, Y. Zhu, and L. Wang, “A complete CPU-FPGA architecture for protein identification with tandem mass spectrometry,” in 2019 International Conference on Field-Programmable Technology (ICFPT). IEEE, 2019, pp. 295–298.

[10] S. Kumar and F. Saeed, “Communication-avoiding micro-architecture to compute Xcorr scores for peptide identification,” in 2021 31st International Conference on Field-Programmable Logic and Applications (FPL). IEEE, 2021, pp. 99–103.

